# Microbiological quality assessment of five common foods sold at different points of sale in Burkina-Faso

**DOI:** 10.1101/2021.09.28.462219

**Authors:** Muller K. A Compaore, Dissinviel Stéphane Kpoda, Bazoin S. R. Bazie, Marcelline Ouedraogo, Alphonse Yakoro, Fulbert Nikiema, Asseto Belemlougri, Romaric Meda, Moumouni Bande, Nicolas Barro, Elie Kabre

## Abstract

The aim of the present study was to assess the microbial quality of five different types of food such as bread, pasta, rice with sauce, beans and milk sold in five localities of Burkina Faso. One hundred and one (101) samples were collected and microbial quality were assessed by evaluating the food hygiene indicators such as total aerobic mesophilic flora, total coliforms, thermotolerant coliforms, yeast and mould. Food safety indicators such as *Escherichia coli, Salmonella*, coagulase-positive staphylococci, *Clostridium perfringens* and *Bacillus cereus* were checked too. All samples were analyzed under ISO methods.

The results showed that 73.27% of samples were satisfactory while 14.85% were acceptable and 11.88% were not satisfactory according to international standards. Among the food safety indicators sought, *Escherichia coli* was detected in two samples and *Bacillus cereus* in four samples. Most of the analyzed food exhibited good hygiene behavior within the acceptable limits and the highest of not satisfactory rate was observed in milk powder and rice with sauce. Ouagadougou samples record the highest number of not satisfactory samples.

Despite the general quality was satisfactory, the presence of specific microorganisms such as coliforms is indicative of the poor hygiene surrounded these foods. It is therefore necessary to train and follow up the vendors in the handling of equipment, hand-washing practices and selling environment hygiene for better improvement of the quality of the street foods.

## 1. Introduction

Food quality is always a concern when intended to human consumption. Food-borne diseases have been increasing in recent years, with a greater impact on the health and economy of developing countries than developed countries [1]. According to World Health Organization (2020) access to sufficient amounts of safe and nutritious food is key to sustaining life and promoting good health and unsafe food containing harmful bacteria, viruses, parasites or chemical substances, causes more than 200 diseases ranging from diarrhea to cancers. An estimated 600 million (almost 1 in 10 people in the world) fall ill after eating contaminated food and 420 000 die every year, including 125 000 children under the age of 5 years [2]. Common foods such as bread, beans, pasta, rice with sauce and powdered milk are common dishes suited as well as to many low-income people as to living conditions in large cities. These foods refer to street foods that play an important role in developing countries such as Burkina Faso. According to the Food and Agriculture Organization, Street foods are ready-to-eat foods and beverages prepared and/or sold by vendors and hawkers especially in streets and other similar public places. This definition emphasizes the retail location on the street, with foods sold from pushcarts, bicycles, baskets or balance poles, or from stalls that do not have four permanent walls [3]. These street foods feed millions of people daily with a wide variety of ready-to-eat foods and beverages sold and sometimes prepared in the street or public places, relatively cheap and easily accessible [4,5]. Although in developing countries the informal food vending sector has in recent years grown into a lucrative trade that competes with the formal sector, ignorance in regard to and inadequate knowledge of food handling practices, together with a lack of formal education, have prevailed among the majority of informal food handlers [6,7]. Food can serve as ideal culture medium for the growth of microorganisms which can cause decomposition, spoilage and also serve as a vehicle for transmission of food borne illness [8]. The food production sites are generally located either in living spaces or near workplaces or on sale sites. The majority of these vending sites lacks basic infrastructure and services such as potable running water and waste disposal facilities, hand and dishwashing water is usually insufficient and often reused, sometimes without soap, waste water is discarded in the street and garbage often disposed of in the vicinity of the stall [6,9].

Street food quality is a concern around the world. Studies conducted in many countries such as Mozambique [10], Malaysia [7,11], Brazil [12], Bangladesh [13], Kenya [14] and so on have been reported. In developing countries, street foods are drove by men and women that knowledge and expertise in food handling are often limited and they often engage in street food mainly to escape poverty, especially as little start-up capital is required [15]. However, it is important to consider the health and safety impact of these food products, because foodborne infections are more and more frequent, hence the need for control strategies to ensure food safety and consumer protection. According to [8] foodborne illness is a major universal health issue in developing countries due to difficulties in safe guarding food from cross-contamination.

Burkina Faso is a land lock country neighbored by six countries, with which it shared almost the same habits of street food accessibility. Vending foods on the street is a common aspect of lifestyle in countries in which there are high unemployment, low salaries, limited work opportunities and limited social programs [16,17]. Foods sanitary quality controls are often limited to large urban centers to the detriment of rural populations. This study is intended to investigate and shed light on the microbial safety of five commons street foods sold at different points of sale in five localities of Burkina-Faso.

## 2. Material and methods

### 2.1. Sample collection and storage

One hundred and one samples divided into five groups including 15 bread samples, 12 for beans samples, 12 pasta samples, 19 rice with sauce samples and 43 milk powder samples were collected in five localities of Burkina Faso.

Table I shows the repartition of all the food samples submitted to this study.

**Table I:** Repartition of the food samples

| Samples | Ouagadougou | Bobo-Dioulasso | Dakola | Cinkansé | Niangoloko | Quantity | Percentage % |
| --- | --- | --- | --- | --- | --- | --- | --- |
| Beans | 4 | 2 | 2 | 2 | 2 | 12 | 11,88 |
| Milk powder | 15 | 9 | 6 | 6 | 7 | 43 | 42,57 |
| Bread | 4 | 3 | 2 | 3 | 3 | 15 | 14,85 |
| Rice with sauce | 6 | 4 | 3 | 3 | 3 | 19 | 18,81 |
| Pasta | 3 | 3 | 2 | 2 | 2 | 12 | 11,88 |
| Total | 32 | 21 | 15 | 16 | 17 | 101 | 100 |

Sampling sites include ordinary restaurants, local markets and shops. The concerned localities were Ouagadougou, Bobo-Dioulasso, Dakola, Cinkansé and Niangoloko. All samples were taken aseptically in sterile plastic bag, kept in an insulated cold box containing ice boxes or stored at room temperatures. Samples reached the laboratory are immediately analyzed or kept under 4°C until used. Figure 1 indicated the sampling localities.

**Figure 1:**
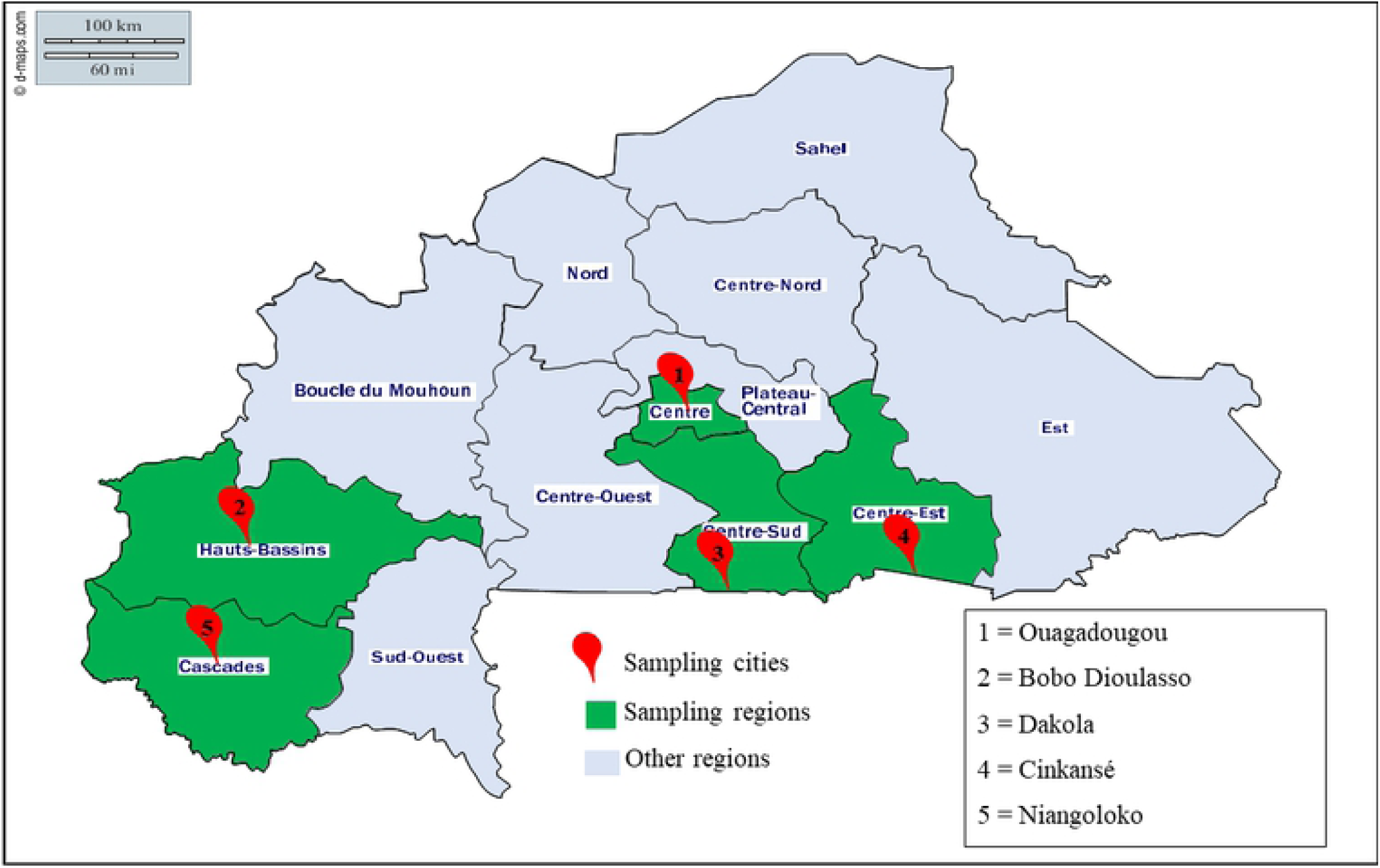
Source: Adapted from https://www.d-maps.com/carte.php?num_car=25733&lang=fr

### 2.2. Analysis parameters

Parameters applied to each group of samples are those recommended by the *Codex Alimentarius*. In addition, Anaerobic Sulfito Reductive (ASR) bacteria, *Clostridium perfringens* and *Bacillus cereus* were investigated in milk only.

#### Microbial Analysis

twenty-five (25) grams of each food sample were homogenized into 225 mL of sterile buffered peptone water (Liofilchem diagnostic, Italy) and homogenized in a Bag Mixer (Interscience, France). Further tenfold serial dilutions were made with in sterile buffered peptone water. Duplicate plates were made for each sample at each dilution under ISO 6887 standard methods. Microbial counts were expressed as colony-forming units per gram (CFU/g).

### 2.3. Evaluation of the food hygiene indicators

#### Total aerobic mesophilic flora

were counted among all the food samples onto standard plate count (PCA) agar (Conda Pronadisa, Spain) under **NF ISO 4833: 2003**. Plates were incubated at 30 ± 1°C for 72 h. After incubation the number of colonies was counted on the plate with less than 300 colonies.

#### Coliforms

that are known to be an indicator of fecal contamination were counted onto standard violet red bile lactose (VRBL) agar (Conda Pronadisa, Spain) and incubated at 37°C for 24 to 48 hours under ISO 4832:1991. Only the Petri dish containing less than 150 colonies were considered.

#### Thermotolerant coliforms

also known to be an indicator of fecal contamination were counted onto standard violet red bile lactose (VRBL) agar (Conda Pronadisa, Spain) and incubated at 44.5 ± 0.5°C for 24 to 48 hours under **NF V08-017:1980**. Only the Petri dish containing less than 150 colonies were considered.

#### Yeast and mould

were counted onto standard yeast extract glucose chloramphenicol (YGC) agar (HiMedia Laboaratories, India) and incubated at 25 ± 1°C for 5 days following **ISO 7954:1988**. The growth of moulds was checked every day in order to avoid invading colonies. Considered Petri dish for bacterial counting were less than 150 colonies.

### 2.4. Evaluation of food safety indicators

#### Escherichia coli

*E. coli* were identified through the IMViC test from thermotolerant coliforms. Briefly, suspected colonies from thermotolerant coliforms were selected and subcultured on Nutrient Agar at 37°C for 24 hours. Pure cultures grown on Nutrient Agar were used for Oxidase test and determination of IMViC pattern (indole, methyl red, Voges Proskauer and citrate utilization test) under the Standard Procedures for food Analysis. Positives clones were transferred into Levine BBLTM Eosin Methylene Blue Agar (EMB) Agar France, which was incubated at 37 ± 1°C for 24 hours. *Escherichia coli* ATCC 8739 was uses as positive control for all analyses.

#### Salmonella spp

*Salmonella* species were investigated according to the standard - Horizontal method for detection of *Salmonella spp* **ISO 6579:2007**. Briefly, the non-selective enrichment was done by adding 25 g of each sesame sample into 225 mL buffered peptone water (Liofilchem diagnostic, Italy) and homogenized in a Bag Mixer (Interscience, France). Incubation was done at 37°C for 18 to 20 hours. The selective enrichment step was performed onto both tetrathionate (Müller-Kauffman) (Liofilchem diagnostic, Italy) and Rappaport Vassiliadis Soy (Difco laboratories) broths incubated respectively at 37 ± 1°C and 42 ± 1°C for 18 to 20 hours. A brilliant green at 0.95% was added to the selective media Tetrathionate broth in order to inhibit the growth of Gram-positive bacteria. Selective isolations were performed onto Xylose Lysine Deoxycholate (HiMedia Laboaratories, India) and *Salmonella-Shigella* (HiMedia Laboaratories, India) agars. Suspected colonies were purified on nutrient agar and then submitted to API 20E (BioMérieux, France) test for biochemical confirmation. *Salmonella typhimurium* (ATCC 14028) and *Salmonella enteritidis* (ATCC 13076) was used as positive control. The Key biochemical tests including the fermentation of glucose, negative urease reaction, lysine decarboxylase, negative indole test, H2S production, and fermentation of dulcitol [18].

#### Coagulase-positive staphylococci

Staphylococci were isolated under **ISO 6888-1:1999**. Briefly, 25 g of each food sample were dissolved in 225 ml of Pepton Water (Liofilchem diagnostic, Italy). and homogenized in a Bag Mixer (Interscience, France). A loop of 0.1 ml of each sample were then seeded onto Baird Parker Agar (BP) supplemented with egg-yolk tellurite emulsion (HIMEDIA) and incubated under aerobic conditions at 37 °C for 24 and 48 h. The samples producing typical colonies (grey-black, surrounded by a dull halo) were considered. Biochemical confirmation to determine whenever these colonies are Coagulase positive was performed using rabbit lyophilized plasma.

#### Anaerobic Sulfito Reductive (ASR) bacteria and Clostridium perfringens

ASR were isolated under **ISO 15213:2003**. For this purpose, 25 g of each food sample were dissolved in 225 ml of Peptone Water (Liofilchem diagnostic, Italy). and homogenized in a Bag Mixer (Interscience, France). 1 ml from each dilution were mixed with tryptose sulfite cycloserine agar and after solidification the plates were overlayed by using the same medium. After incubations at 46°C for 18 to 20 h under anaerobic condition, characteristic colonies were isolated for biochemical confirmation of *Clostridium perfringens* under **ISO 7935:1997**. Briefly, five black colonies were picked, and each was inoculated into 10 ml of fluid thioglycolate broth. After 18 to 20 h at 37°C, thioglycolate tubes were used to inoculated complete lactose sulfite broth containing Durham tubes and then incubated at 37°C for 18 to 20 h. Tubes with black butt and gas in the Durham tubes are considered as *Clostridium perfringens. Clostridium perfringens* ATCC 13124 was used as a positive control for all biochemicals tests.

#### Bacillus cereus

Enumeration of *Bacillus cereus* was performed by surface plating techniques of 100µl on mannitol-egg yolk-polymyxin under ISO 7932:1993. This media use polymyxin B as the selective agent and permit presumptive identification by the lecithinase reaction on the egg yolk and the inability of *Bacillus cereus* to catabolize mannitol. The media were incubated at 35 °C for 24 h. The number of *Bacillus cereus* was determined after enumeration of the colonies having its characteristic appearance and submitted to biochemical characterization. *Bacillus cereus* ATCC 11778 was used as a positive control.

The appreciation criteria of microbiological quality of all food samples are enumerated in table II

**Table II:** *Appreciation criteria of ready to eat meals (A) and conditioned milk powder (B);* Source:[19]

|  | Parameters | Criteria m UFC/g | M (3m) UFC/g |
| --- | --- | --- | --- |
| Ready meals | *Salmonella* | Absence/25g | ---- |
|  | Staphylococci | 10<sup>2</sup>/g | 3x10<sup>2</sup>/g |
|  | Total aerobic mesophilic flora | 3x10<sup>3</sup>/g | 9x10<sup>3</sup>/g |
|  | Coliforms | 10<sup>3</sup>/g | 3x10<sup>3</sup>/g |
|  | Thermotolerants Coliforms | 10/g | 3x10/g |
|  | *E. coli* | 10/g | 3x10/g |
|  | Yeast and Mould | 10<sup>4</sup>/g | 3x10<sup>4</sup>/g |

B
|  | Parameters | Criteria m UFC/g | M (3m) UFC/g |
| --- | --- | --- | --- |
| **Conditioned milk powder** | Salmonella | Absence/25g | ---- |
|  | *Staphylococcus aureus* | 10/g | 30/g |
|  | Total aerobic mesophilic flora | 5x10<sup>4</sup>/g | 1.5x10<sup>5</sup>/g |
|  | Coliforms | 1/g | 3/g |
|  | Thermotolerants Coliforms | Absence | ---- |
|  | *E. coli* | Absence | ---- |
|  | Yeast and Mould | Absence | ---- |
|  | ASR | 10/g | 30/g |
|  | *Clostridium perfringens* | 1/g | 3/g |
|  | *Bacillus cereus* | Absence | ---- |

### 2.5. Statistical analysis

Statistical analysis was done using IBM SPSS Statistics Version 20.0.0. and graphs were built under Microsoft Excel 2016. The mean value and standard deviation were calculated from the obtained data in each food sample group. Comparison test was done to determine whenever there were significant differences (*P* ≤ 0,05) within these types of food samples.

## 3. Results

The results showed that out of 101 samples submitted to this study, 73.27% were satisfactory according to the criteria used, while 14.85% were acceptable and 11.88% were not satisfactory (Figure 2). Table III summarizes the raw results of the evaluation of the bacterial load and the presence of pathogens in all the samples submitted to this study. Only 6.93% (7 samples over 101) of the analyzed samples did not show any microorganism. The highest number of coliforms, total aerobic mesophilic flora, thermotolerant coliforms and yeast and mould were recorded in the same sample of rice with sauce in Ouagadougou and values are respectively 2×10^4^, 2.4×10^5^, 1.7×10^4^ and 1.6×10^4^. The pathogens *E. coli* were found in two (02) samples of Ouagadougou namely bean and pasta. On the other hand, *Bacillus cereus* was found in four (04) samples of milk namely two in Ouagadougou and two in Cinkansé. Neither *Salmonella* nor *Clostridium perfringens* were found in any samples.

**Figure 2:**
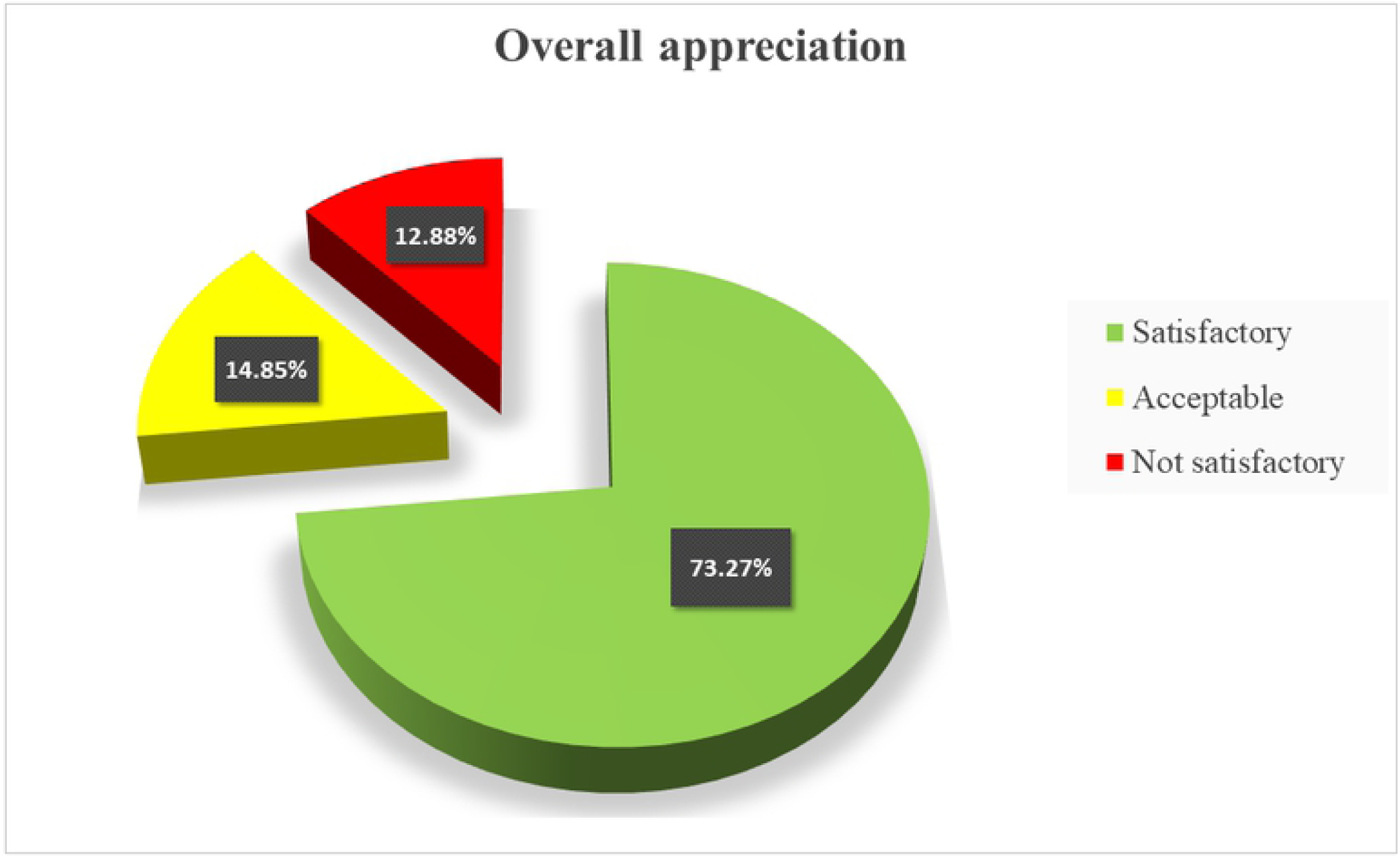
the overall appreciation of samples quality.

**Table III:**
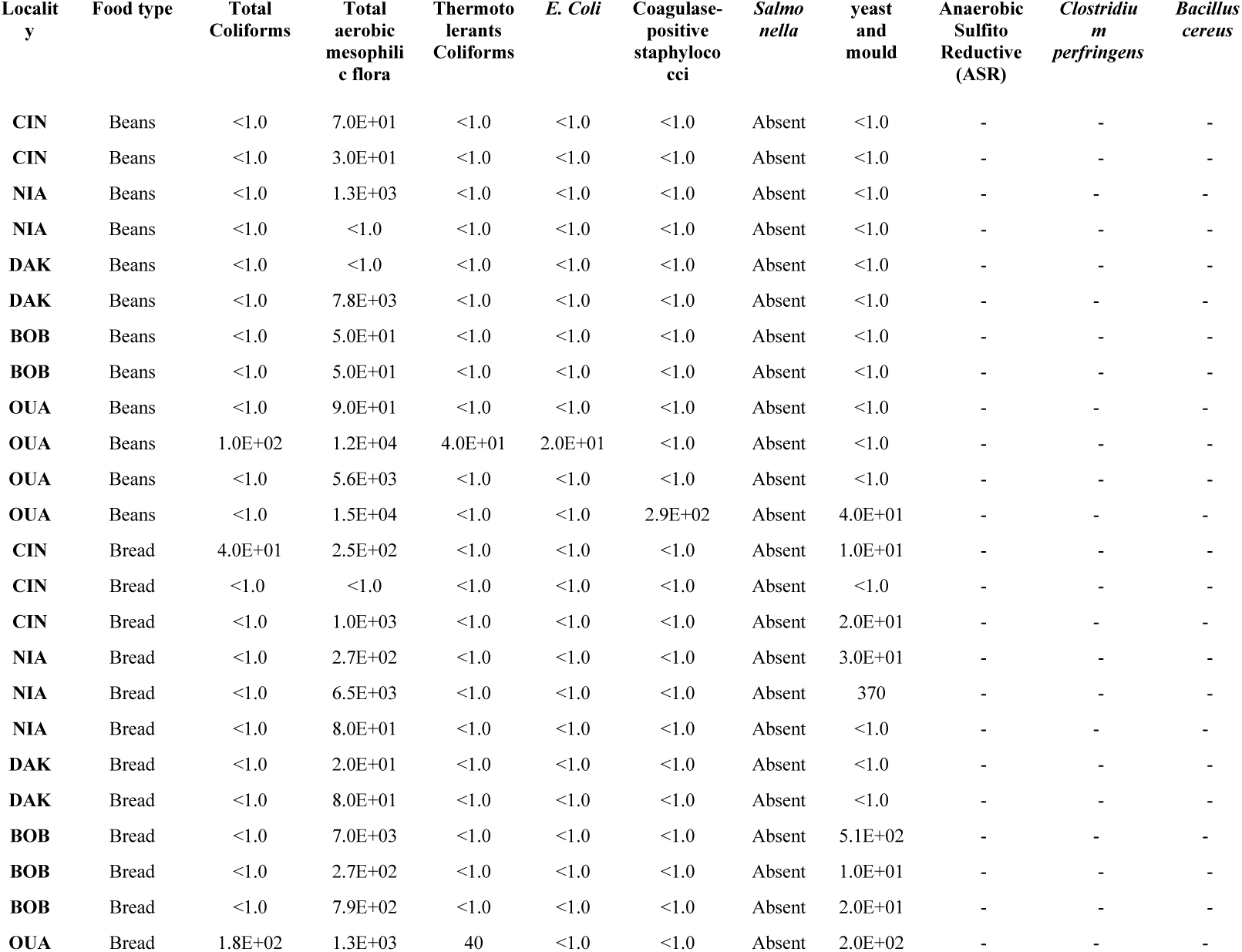
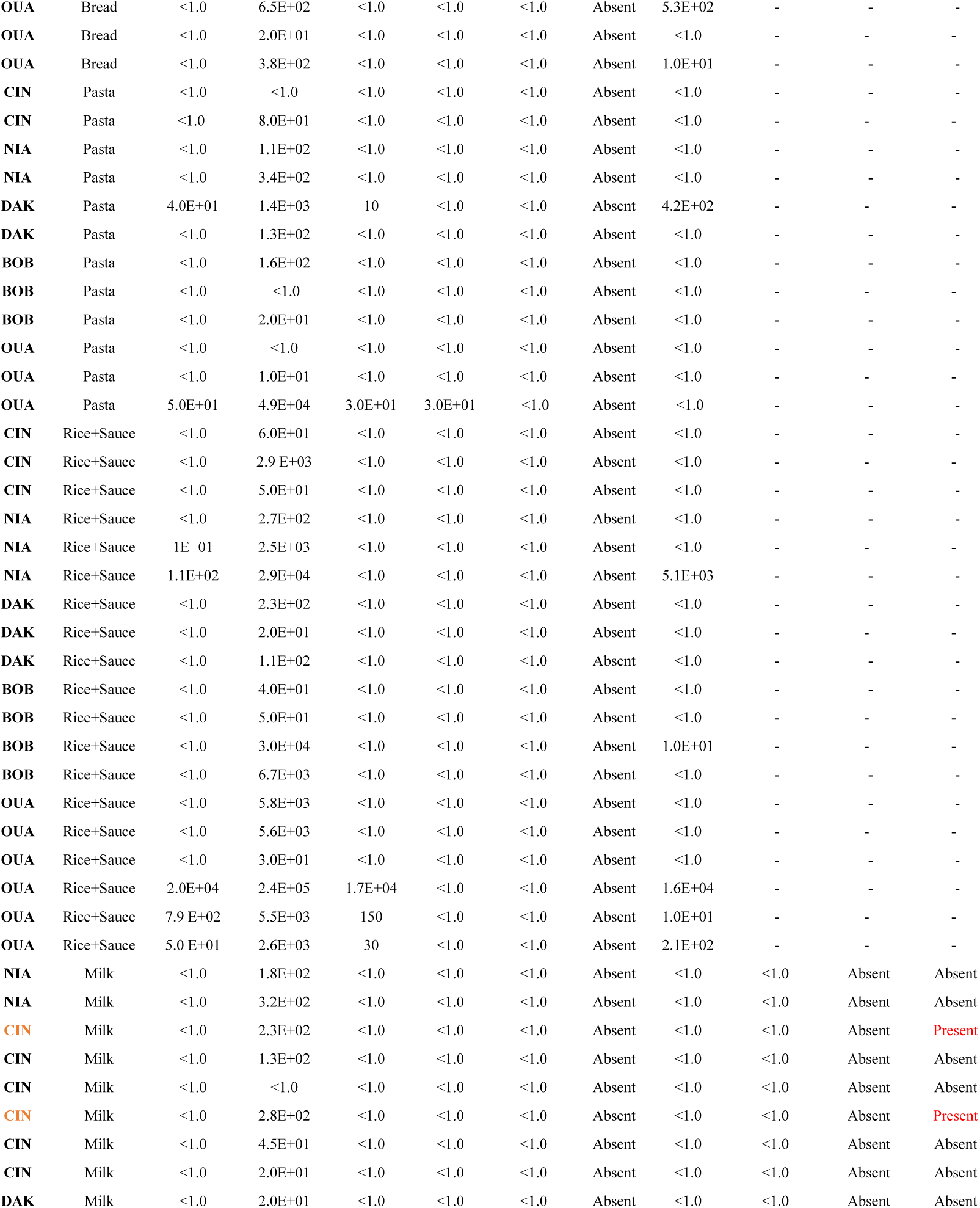
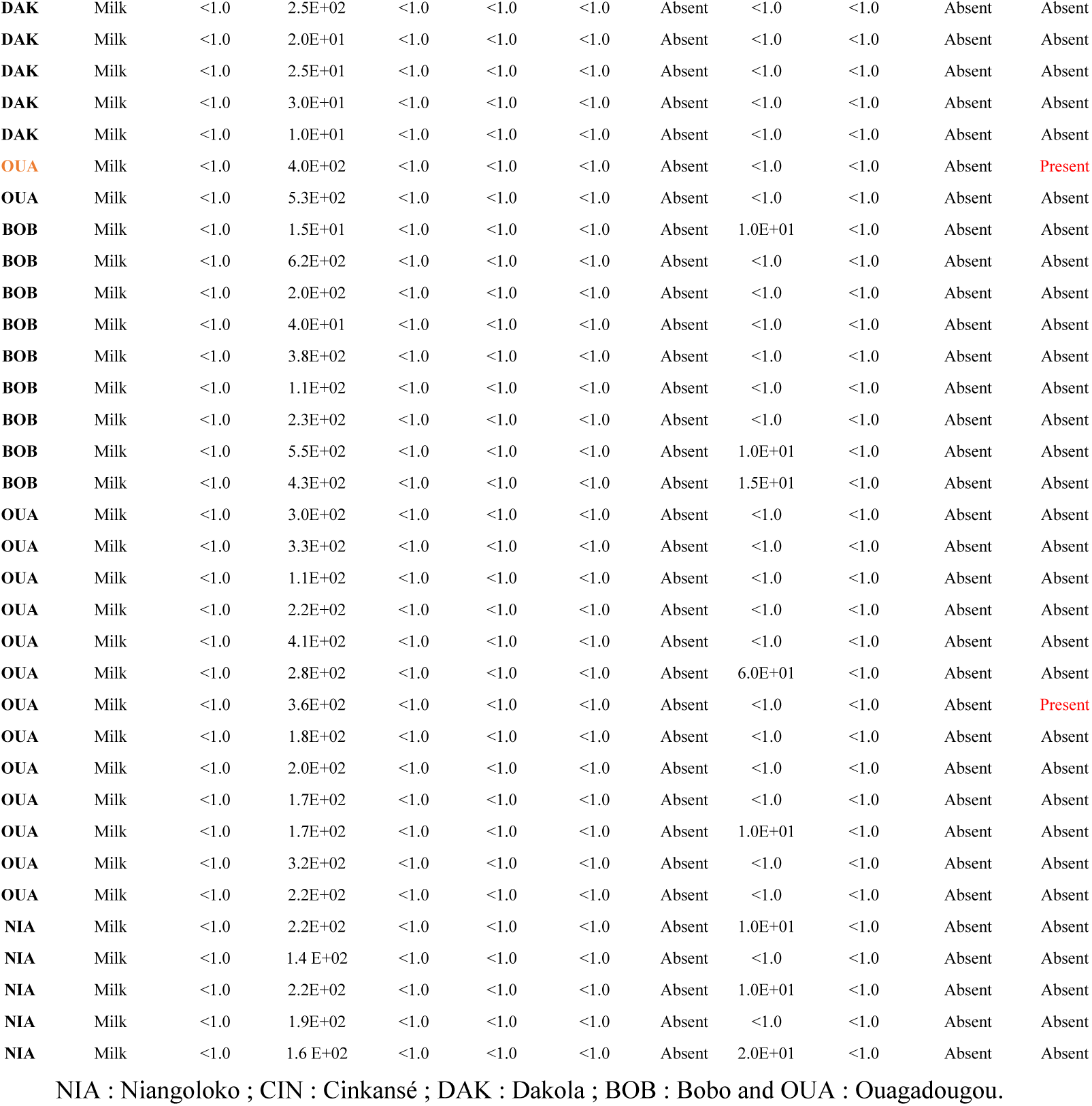
Microbiological analyses

Figure 3 summarizes the overall appreciation of all samples according to cities. Ouagadougou has the highest number of samples and the highest number of not satisfactory (7.9%). Off set Dakola, all the other localities showed at least sample one that is not satisfactory.

**Figure 3:**
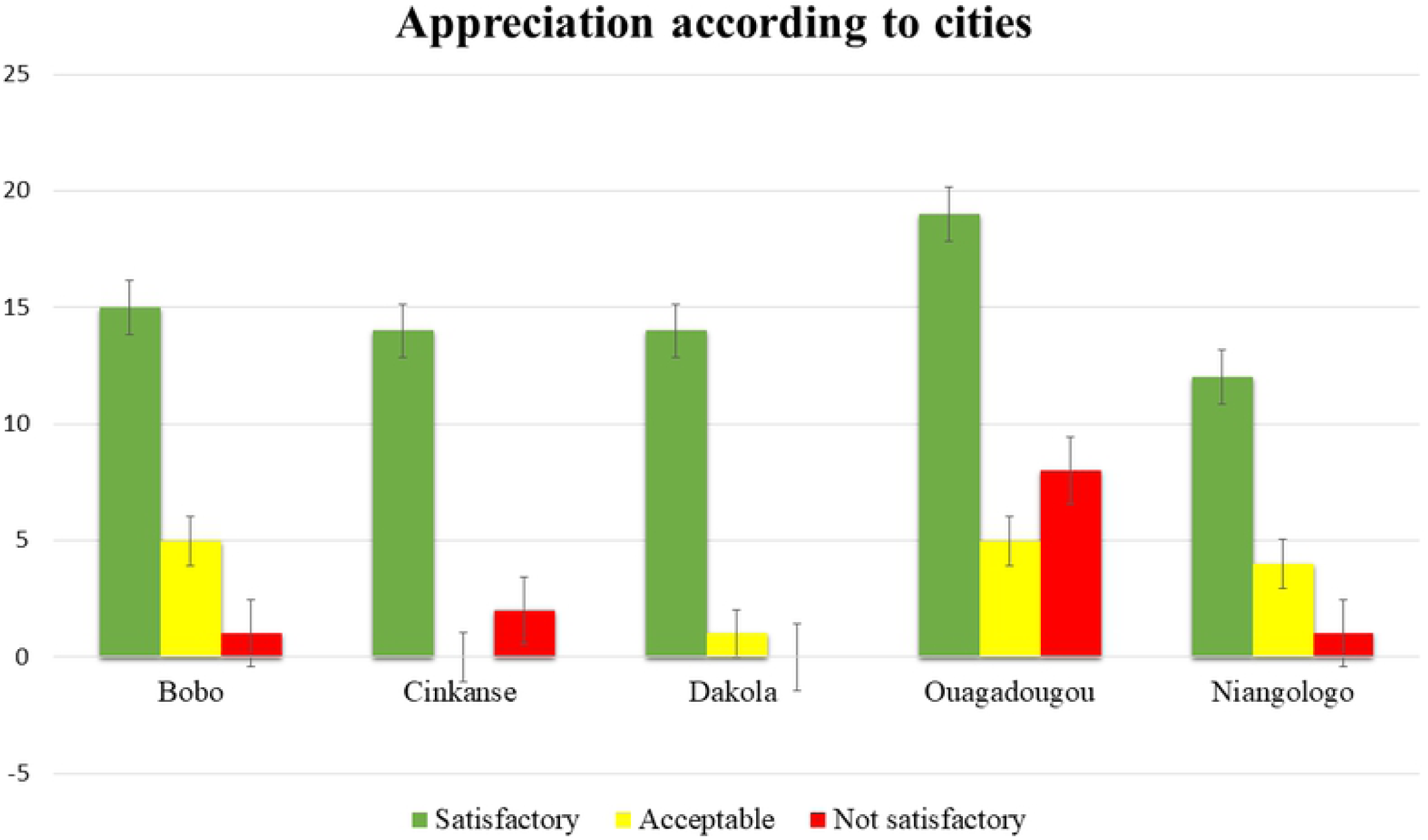
Appreciation according to cities.

The figure 4 gives an overview of the appreciation of the samples according to their nature. The milk samples represent the highest percentage i.e., 42.57%. All samples showed at least one case that is not satisfactory. Milk powder and rice with sauce record the highest cases of not satisfactory at equal value, i.e., 3.96%.

**Figure 4:**
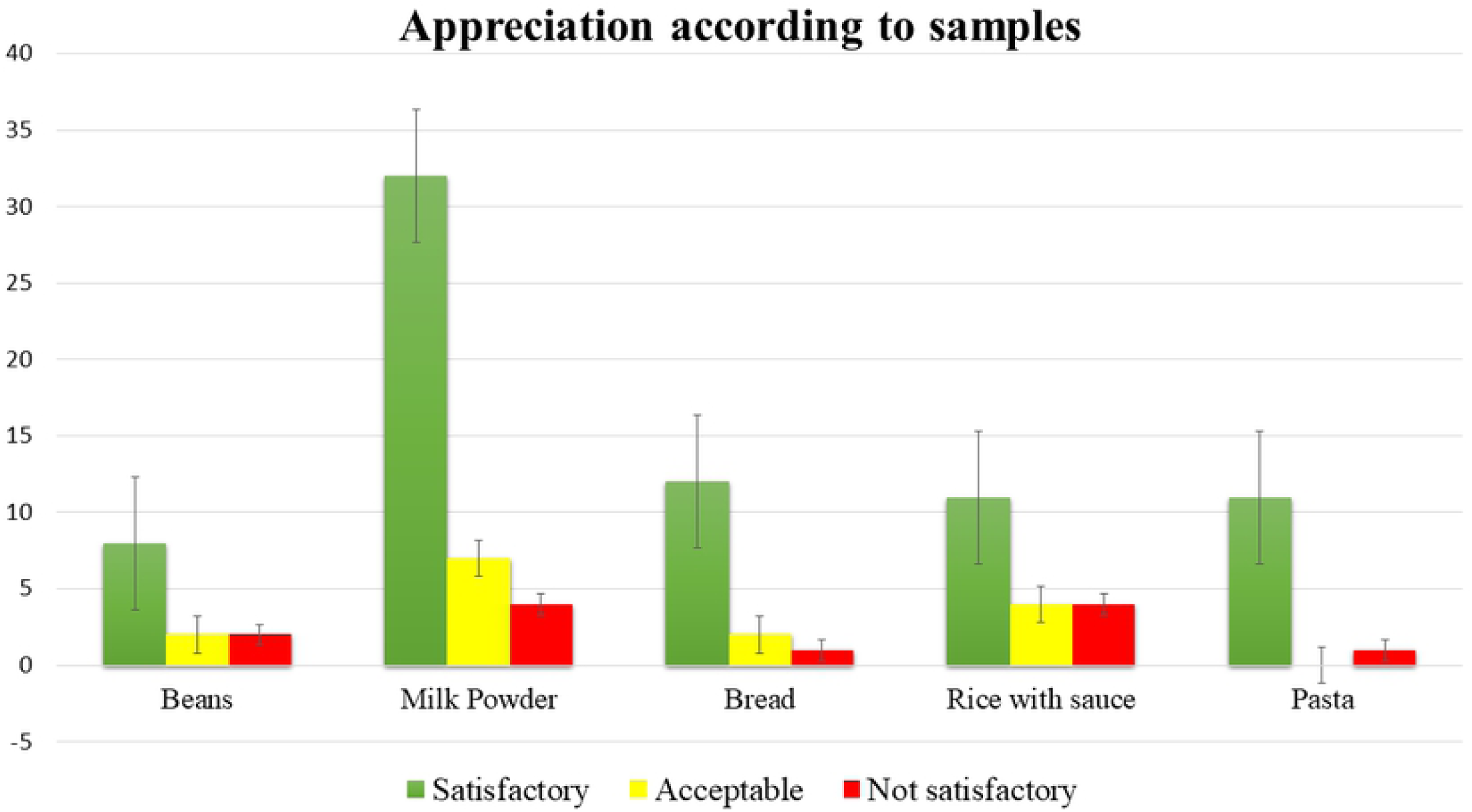
Appreciation according to samples.

Depending on the profile of the microorganism sought, the total aerobic mesophilic flora was present in 93.07% of all samples, yeast and mould in 24.75%, total coliforms in 9.90%, thermotolerant coliforms in 6.93%, *E. Coli* in 1.98% and coagulase-positive staphylococci in 0.99%. *Bacillus cereus* was present in 9.30% of milk sample only. Neither *Salmonella* and anaerobic sulfito reductive (ASR) nor *Clostridium perfringens* were detected in any samples.

## 4. Discussion

The control of foodborne pathogens is an essential measure in preventing the appearance and spread of foodborne diseases in the population [20]. According to FAO (2009), unsafe food poses global health threats, endangering everyone; infants, young children, pregnant women [2]. Foods submitted to this study feed millions of low-income persons daily and therefore must be of very good nutritional and microbiological quality.

Despite the relatively poor sale environment hygiene noticed on the field; these foods were in general, microbiologically safe as only 11.88% were not satisfactory according to the standards used. Similar satisfactory results of street food were reported with breakfast and snack foods in Ghana [4], with meat and chicken stews and maizemeal porridge.in South Africa [21]. Most of the unacceptable limits derived from the total viable counts where microorganisms were present in 93,07% of samples, independently of their pathogenicity and the presence of coliforms. Total viable numbers, might come from all sources surrounding the foods, mostly from air helped by wind. The highest total viable counts recorded was 2.4×10^5^ and was found in rice with sauce. Published papers report high total viable counts in street foods from 10^5^ up to 10^9^ UFC/g [22,13,23]. [24] report that many foods provide an environment conducive to microbial growth, and indicator counts in such foods may reflect the time and conditions of storage. Otherwise, total viable counts cannot be used as a safety indicator, as there is generally no correlation between its value and the presence of pathogens or their toxins [24]. Excepted milk samples, all foods samples submitted to our study undergo heating step that should normally reduce the microorganism population. That should be one of the reasons, we found out low total viable counts as compared to other studies. Microorganisms detected seems to have post contamination origin due to food and material handling. Street foods contamination mainly occur through hands [5,1,11,25]. Moreover, the presence of indicator bacteria in ready-to-eat food, although not inherently a hazard, can be indicative of poor practice that may be poor quality of raw materials or food components; undercooking; cross-contamination; poor cleaning and poor temperature and time control [26]. The presence of thermotolerants coliforms and total coliforms respectively at 6,93% and 9,90%, in the study samples confirm these finding of post-processing contamination. The thermotolerant coliforms have the same properties as the total coliforms at the difference that lactose fermentation occurred at 44.5°C ± 0.5. However, Coliforms are known to be a possible fecal contamination indicator. That suggest the possibility of potential presence of other enteric bacteria that can be pathogens. *E. coli* were detected in 1.98% of study samples specifically in bean and paste samples from Ouagadougou city. Their presence in heat treated foods might therefore signifies inadequate cooking or post-processing contamination. According to [6], samples collected during the holding period after the food had already been exposed to high temperature processing, any presence of *E. coli* could only be attributed to faecal contamination from the hands of food handlers and/or from contaminated working surfaces and utensils. Therefore, special attention should be paid to the street food chain with reference to the 5Ms (Main œuvre, Matière, Milieu, Matériels et Méthode) as decreed in the *Codex Alimentarius*. In addition, all food handlers must pay particular attention to their personal hygiene and hand washing after handling raw food and after using the toilet.’

Coagulase-positive staphylococci were detected in a bean sample of Ouagadougou at the level of 2.6×10^2^. The main reservoir of staphylococci in humans is the nostrils, although staphylococci can also be found on hand [1]. One might think that Coagulase-positive staphylococci contamination of our bean sample comes from handling, perhaps because of more frequent hand contact during preparation and serving. Staphylococcal food poisoning is one of the most common food-borne diseases in the world following the ingestion of staphylococcal enterotoxins that are produced by enterotoxigenic strains of coagulase-positive staphylococci, mainly Staphylococcus aureus [27]. Coagulase-positive staphylococci have been found in a large number of commercial foods by a wide range of investigators [6,27,22,23,8,14] but their appear to be more present with high numeration as compared to our results.

Yeasts and molds are commonly enumerated in foods as quality indicators and they have no predictive value for the occurrence of toxigenic fungi or other pathogens[24]. They are responsible of food spoilage when these foods are exposed to ambient condition without any protection. This study exhibits 24.75% of yeast an molds contamination that load varying from 1.0×10 to 1.6×10^4^ UFC/g. The presence of yeasts and molds in heat-treated foods might also has its roots in inadequate cooking, post-processing contamination, cross-contamination or even poor quality of raw materials. [28] find out similar results on yeast load varying from 1.2 up to 5.2 logUFC/g while assessing the microbiological quality of ethnic street foods in the Himalayas.

Control parameters applied to milk samples differ from those applied to the others samples as Anaerobic Sulfito-Reductive (ASR), *Clostridium perfringens* and *Bacillus cereus* were also searched. Large numbers of *Bacillus cereus* are needed to cause illness either by releasing toxin into the food prior to consumption (emetic syndrome) or by producing a different toxin or toxins in the gut after eating the food (diarrheal syndrome) [26]. *Bacillus cereus* was isolated from only four (4) milk powder samples (9,30%). Milk powder samples submitted to this study were somehow reconditioned by the vendors. The predominance of *Bacillus cereus* was probably due to cross contamination of bacillus spores present at the conditioning environment. Similar results were obtained in South Africa where nine (9) meat/chicken samples (10.3%) and 6 maizemeal porridge samples (5.3%) were positive for *Bacillus cereus* [21].

Dakola, Cinkansé and Niangologo localities, which are the border post cities with high traffic and where food handlers lack of hygiene facilities (water supply, expertise…) like Ouagadougou and Bobo, were expected to have a high contamination load. Surprisingly, Ouagadougou recorded the highest rates of not satisfactory sample followed by Cinkansé. It can be hypothesized that this might be due to the high number of Ouagadougou sampling (31.68%) as compared to the other samples.

The food safety indicators such as *Salmonella*, the Anaerobic Sulfito Reductive (ASR) and *Clostridium perfringens* were not detected in any of all samples. Similar results were found in South Africa while trying to determine the health risks associated with street food vending [21]. It appears that important hygiene measures are practiced by almost all food handlers and this is very encouraging. Furthermore, the overall microbiological quality and safety of foods submitted to this research study were within the acceptable limits. One might believe that the advent of covid-19 that has profoundly destabilized developing countries people and fundamentally changed their habits and behaviors might have contributed to reduce the contamination of food through handling. Indeed, the media fuss around handwashing with soap and the use of hydro-alcoholic gels have been accepted by Burkinabe around the country. One of this positive impact might be the reduction of microbial contamination through the hands.

The presence of the different type of microorganisms and the not satisfactoriness samples observed suggested a post contamination during food handling and the possible microbial attacks propagated from the surrounding environment. It is well known that infections caused by microorganisms can be reduced by maintaining correct hand hygiene. Thus, training and education can improve the knowledge of street food handlers and play an important role in risk mitigation. Specific attention should be given for storage and packing processes specific to each food is also required, and the control of water used for utensils and hands washing. That might be one of the best ways to assure constantly a good hygienic quality of street foods.

## 5. Conclusion

This research work highlighted that street food vendors of the study regions of Burkina Faso were able to produce relatively safe foods with low percentage of not satisfactory samples. A void that was necessary was filled, namely the quality of the meals that the average Burkinabe consumes every day, especially in border areas and urban centers. The microbial quality of these foods was acceptable in general even some fecal indicators are still presents suggesting the possibility of potential presence of other enteric bacteria that can be pathogens. As the different part of the country share the same street foods habit, one could extrapolate those similar behavioral patterns may be found elsewhere within the whole country. Therefore, education and training of foods handlers is crucial to control potential food borne illness. One might believe that this study is the first of a series of several street foods quality control to prevent intoxication outbreak that can impact negatively the economy of developing countries that is already weak.

## Acknowledgements

This work was initiated by the PAASME-UE project (“Productions et analyses des données pour améliorer la santé de la mère et de l’enfant au Burkina Faso”) in collaboration with the INSP, financed by the European Union. At the end of this work, it is important to thank:

- The National Institute of Public Health (INSP) for the scientific collaboration;
- The European Union (EU) for the financing of this project;
- The Health Research Ethics Committee for authorizing the conduct of this study;
- The Regional Health Directorates of the Centre, Hauts-Bassins, Cascades and Centre-East regions for authorizing the conduct of this study;
- The populations of the Centre, Hauts-Bassins, Cascades, Centre-East and Centre-South regions for their collaboration.

